# Aeroplane wing, a new recessive autosomal phenotypic marker in malaria vector, *Anopheles stephensi* Liston

**DOI:** 10.1101/2023.06.05.543679

**Authors:** Chaitali Ghosh, M Soumya, Naveen Kumar, R Chethan Kumar, Soumya Gopal Joshi, Sampath Kumar, Suresh Subramani, Sunita Swain

## Abstract

A novel and distinct mutant, with an aeroplane wing (*ae*) phenotype, is reported for the first time in the urban malaria vector, *Anopheles stephensi*. The mutant mosquitoes exhibit extended wings that are easily visible to naked eyes in both sexes. This mutant was first observed in a nutritionally stressed experimental isofemale line and characterized for its genetic inheritance and other related parameters. Meticulous and strategic genetic crosses revealed that the *<u>ae</u>* gene is an autosomal, recessive, non-sex-linked and monogenic trait with full penetrance and uniform expression in the adult stage. Cytogenetic study of the ovarian polytene chromosome revealed an inversion on the 3L chromosome (3L*i*) in both the *ae* mutant and its parent line. No significant differences in wing venation and other parameters were observed in *ae* mutants compared to their normal parental lines. This *ae* mutant would be an excellent marker that can be used by researchers to study the function of related genes within the genome.

**Author summary:** In the present study, we have established and describe the inheritance of an unusual novel aeroplane (*ae*) winged mutant in *Anopheles stephensi*, an urban malaria vector in India. The *ae* mutant lines exhibit three open-wing orientations in both the sexes of adults viz. left (LW), right (RW) and double wing (DW) during its resting phase. Through various inbreeding crosses we demonstrated the mode of inheritance of the *<u>ae</u>* gene to be autosomal, recessive and monogenic in nature. The morphometric studies of eggs and wings revealed that mutant lines are on par with their parental lines. Cytogenetic study of polytene chromosome of *ae* mutant revealed the presence of heterogenic inversion on the 3L chromosome arm, which might help in adaptation mechanism. We strongly believe that *ae* phenotypic markers have great applications bridging both basic and applied genetic research such as constructing linkage maps, identifying loci of quantitative and/or qualitative traits and as guides for insect transformation studies.

## Introduction

Mutant phenotypic markers are well described in model organisms and have played an important role in characterizing their genomic information. These mutations have helped geneticists in designing many genetic crossing experiments to understand the inheritance pattern that can be used in translational research. The first phenotypic genetic marker in the fruit fly, *Drosophila melanogaster,* was established through experimental mutagenesis that allowed T. Morgan to establish the first genetic maps of *Drosophila* chromosomes [1]. Phenotypic markers are visual indicators of characters, such as colour, shape, size, and such observations, dating back over 100 years, aided the discovery of the arrangement and linkage of genes [2, 3]. Even to this day, morphological markers are very useful tools in understanding genetics, breeding practices and act as a vital bridge between classical and molecular genetics. Though several such milestones have been achieved in model organisms, other insect vectors and pests have largely remained elusive. The development and use of various types of markers in such other insect pests and vectors will facilitate faster scientific progress in fundamental and applied research.

Vector-borne diseases have become one of the major public health problems in tropical and subtropical countries including India. *An. stephensi*, Liston, 1901 is a potential malaria vector in Southeast Asia and the Arabian Peninsula. In the last decade, this species has invaded Africa and Sri Lanka and seems to be spreading, given new reports of its detection [4-6]. Mathematical modelling suggests that over 126 million people may be at risk for *An. stephensi*- transmitted malaria across Africa [7]. *An*. *stephensi* has three biological variants: type, *mysorensis* and intermediate based on the number of egg ridges on egg floats [8]. Such biological forms result from evolution for local adaptation. The process of evolution depends on the occurrence of new mutations that provide genetic variation and influence phenotypic traits by altering gene activity or protein function. Most of these mutations are recognized because of the changes in the phenotype. Over time, as generations of individuals with the trait continue, the advantageous traits become common and establish in the population. The present study was aimed towards the isolation and establishment of *ae* mutant, its phenotypic characterization, and the mode of inheritance of the *<u>ae</u>* mutant gene in *An. stephensi*.

Though limited, there are few reports of such mutant forms in mosquito vectors occurring spontaneously at the larval and adult stages of *An. stephensi*. These include stripe [9] and hairless antenna in adults [10]. Apart from these, eye-colour mutants have also been reported in *An. stephensi*, such as rosy [11], maroon [12], chestnut [13], ruby [14], and yellow body larvae [15]. Tests for allelism were also reported using different larval colour mutants, such as grey and greenish black in *An. stephensi* [16]. Linkage studies have been reported in crosses of the colour mutant green thorax (*<u>gt</u>*) with ruby eye (*<u>ru</u>*) [17]. In the present study, we describe the isolation and genetic analysis of a novel aeroplane-winged (*<u>ae</u>*) mutant in *An. stephensi*. The uniqueness of this mutant line is the visually distinct phenotype of open wings while resting. We have characterized the genetic basis of the *<u>ae</u>* genes by crossing experiments.

The aeroplane (*ae*) mutant has been isolated from an intermediate variant during nutrition challenge experiments under laboratory conditions. Further, the cytogenetic studies on this mutant line have provided valuable insights on inversions and vector evolution.

## Materials and Methods

### A. An. stephensi strains

Genetic crossings, morphometric and cytogenetic studies of *ae* mutant were carried out in the TIGS (Tata Institute for Genetics and Society) insectary. TIGS-2 (T_2_), TIGS-6 (T_6_) and *ae* mutant strains were used in this study. The genetic crosses were performed between T_2_ (w) and mutant (*ae*) lines for characterizing the genetic mode of inheritance of *<u>ae</u>* gene. The parental T_6_ isofemale line was used for comparison. The larvae and adults are reared in the insectary as per the protocol described earlier [18]. Temperature was maintained at 27±1°C, relative humidity (RH) at 75±5% and photoperiod of 12h:12h (L:D).

### (1)#Wild strains

#### TIGS-2 (T_2_)

T_2_ strain was collected from Anna Nagar, Chennai (13.018410°N, 80.223068°E), Tamil Nadu, India in 2016. About 60 gravid females resting on concrete houses were collected with the help of aspirators between 6 and 8 AM. The gravid adults were brought to the insectary and kept inside an adult cage with 10% sugar solution. An ovicup lined with filter paper filling with 1/3-part water was kept for egg laying. The eggs were collected and allowed to hatch. Hatched larvae were transferred to the trays with little amount of larval food. After 9 to 10 days, larvae transformed to pupae, which were transferred to the adult cage for emergence into adults. These adults were used for developing the colony and the strain is being maintained in our insectary for over 60 generations since 2018.

#### TIGS-6 (T_6_)

Similarly, T_6_ strain was collected from Sriramanahalli village (12.972442°N, 77.580643°E), Bangalore rural, Karnataka, India in 2016. About 150 larvae were collected from fresh water in an open cement tank, primarily used for providing water for cattle. The collected larvae along with water from their natural habitat were brought to the insectary. They were allowed to develop into adults and kept inside the cage with 10% sugar solution. They were blood-fed through artificial membrane feeding. The post-blood fed gravid mosquitoes were allowed to lay eggs in an ovicup and followed during filial generations. During the rearing of mosquitoes, the insectary conditions were made to mimic the natural conditions as far as possible (in terms of photoperiods, temperature, humidity, nutrition etc.). This strain is being maintained in the insectary for over 63 generations since 2018.

#### *Development of isofemale line of* T_6_

T_6_ isofemale line was established from a single blood-fed female picked from the T_6_ colony and selected over generations through inbreeding. In every generation, 5 to 10 fully engorged blood-fed females were separated individually in each ovicup for laying eggs. Eggs in each cup are allowed to hatch and maintain separately. The isofemale lines were developed and maintained as per the method published earlier [19].

(2) Mutant strain: Mutant (*ae*) lines were first isolated from a nutrition-deprived stressed colony of the T_6_ isofemale line. The stress experiment was started in the isofemale strain at generation 32. The experiment was designed as follows. In the experimental set, a low amount of larval food (i.e., 0.033±0.001g (33 mg)) was added to the tray containing nearly 100 larvae in 750 ml of RO water along with a control set provided with adequate larval food of 0.230±0.01 g (230 mg, i.e. 7 times more food) with same number of larvae and same amount of water (unpublished data). The *ae* mutants (4 females and 6 males) were noticed the first time in the 5th generation. Mutant adults were collected from each generation and inbred separately. The pure line was established before the crossing experiment was conducted.

### B. Crossbreeding experiment

Genetic crosses were made between the males and females of freshly emerged wild (T_2_) and *ae* mutant adults. To ensure the virginity of male and female adults, individual pupae from both strains were kept in perforated 1.5 ml micro-centrifuge tubes and allowed to emerge into adults, which were screened individually for their sex and released into separate Bugdorm cages (L31 x W31 x H9 cm) for mating. A total of 14 crossing sets were designed for the study [16].

Twenty males and 10 females (2:1 ratio) taken from each line (wild and *ae* mutant) were used for each crossing experiment. Parental crosses were made only between wild types (crosse 1) and pure mutants (cross 2). Reciprocal crosses were made between the parents of pure mutant and wild types (crosses 3 and 4) to generate F_1_ hybrids. Some of the F_1_ hybrids were inbred to obtain F_2_ generation (crosses 13 and 14) and the remaining adults were backcrossed to both parental mutants and wild types. The F_1_ progeny of the crosses 5,6,7, 8 and 13 were derived from the male outcross (cross 3) and the F_1_ progeny of the crosses 9,10,11,12 and 14 were derived from the female outcross (cross 4). Fecundity, % hatchability and number of males and females of both wild and mutant individuals were recorded from every crossing. Chi-square tests were calculated to compare the significance level (*P*>0.05).

Further, F_2_ hybrids were inbred *inter se* to produce F_3_, F_4_ and the process was continued till F_15_ generation. Screening was done to observe the frequency of mutant mosquitoes and scoring of mutants was done from every filial generation.

### C. Morphometric analyses of eggs

A total of 20 individual eggs were randomly selected from mutant colonies. Eggs were placed on a wet filter paper and measured under microscope with an ocular micrometer (Unilab GE- 34, Binocular Research Microscope, India). The measurement of different parameters such as egg shape, size, float length, float width, and float ridge numbers were recorded.

### D. Morphometric analyses of wings

A total of 5 to 6 wing samples from both males and females of *ae* mutants were taken for analyses and wings from their parental T_6_ lines were used as a control for comparison. The wings were carefully dissected on day 10 of eclosion. The mosquitoes were anesthetized on CO_2_ pads and the wings were dissected under a microscope (Olympus, SZX2-ILLK, Germany).

The wings were measured from the distal end of the alula to the tip between R_1_ and R_2_ veins, excluding the fringe scale [20]. Wing images were taken under a stereomicroscope (Leica MZ10F, Germany) and the measurements of wing length and width were recorded using Leica Application Suite X (LAS X) software.

### E. Cytogenetic studies of ae mutant and T_6_ parental line

Cytogenetic studies of polytene chromosomes were carried out from ovarian nurse cells collected from semi-gravid females of the *ae* mutant and parental T_6_ isofemale lines [21]. The semi-gravid females were anaesthetized using CO_2_ pad and placed on a microslide in a drop of modified diluted Carnoy’s fixative solution (Carnoy’s fixative: distilled water, 1:19). The ovaries were dissected and fixed in Carnoy’s fixative (methanol: acetic acid, 3:1) for a few minutes. After fixation, the material was stained with a drop of LAO (lacto-aceto-orcein) for 20 to 25 minutes. After staining, 60% acetic acid was added, and a clean cover slip was placed on the top of the sample. Gentle pressure was applied for squashing. The temporary mounts were sealed with nail polish around the cover slip. The slides were examined for chromosomal inversions under 40X and 60X, respectively, using microscope (Nikon Eclipse, Model SC600). The inversion nomenclature was followed according to the methods of Coluzzi *et al*. [22], Mahmood and Sakai [23] and Sharakhova *et al*. [24].

## Statistical analyses

Descriptive, inferential, and predictive analyses have been carried out in the present study. Significance values were computed using on-line Java Script tests (Chi Square calculator 2×2 - https://www.socscistatistics.com/tests/chisquare/) and for wing measurement calculation was done following t-test analysis for independent or correlated samples using Vassar Stat software (http://vassarstats.net/). The mean and SEM were compared between mutant and parental strains. *P*-value >0.05 was considered statistically non-significant.

## Results

### A. Adults of ae mutants and T_6_ parental strains

Three types of wing orientations were observed in both males and females of *ae* mutant line (Fig. 1). Compared to wild (Fig. 1a and b) mosquito, in left wing (LW) and right wing (RW) mutant, only the left-side (Fig. 1c and d) and right-side (Fig. 1e and f) wings are outstretched respectively, while the other wings remain the same as in wild type. In double wing (DW) mutant, both the left and right wings are widely extended on both sides, forming an angle to the longitudinal axis of the body (Fig. 1g and h).

**Figure 1:**
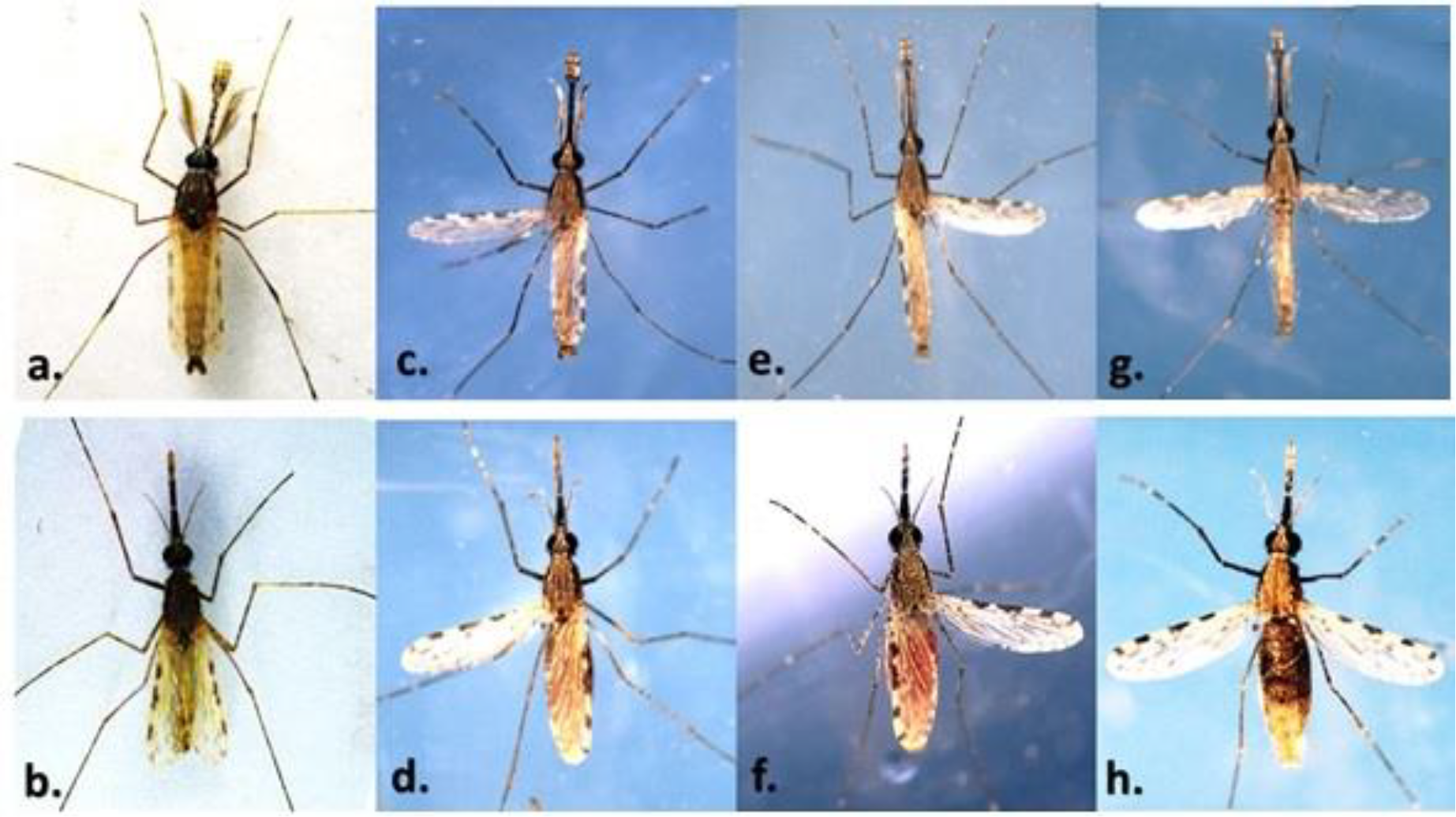
Variation of wing position in *ae* mutant. (a, b) Wild male and female mosquitoes. (c, d) LW male and female mutants. (e, f) RW male and female mutants. (g, h) DW male and female mutants.

### B. Inheritance of <u>ae</u> mutant gene

Testing the mode of inheritance of *<u>ae</u>* gene involved 14 crosses between the *ae* mutants and wild types. Progeny derived from each cross were analysed. Schematic illustration of different crosses is presented in Fig. 2. Results of these crosses are summarized in Table 1.

**Figure 2:**
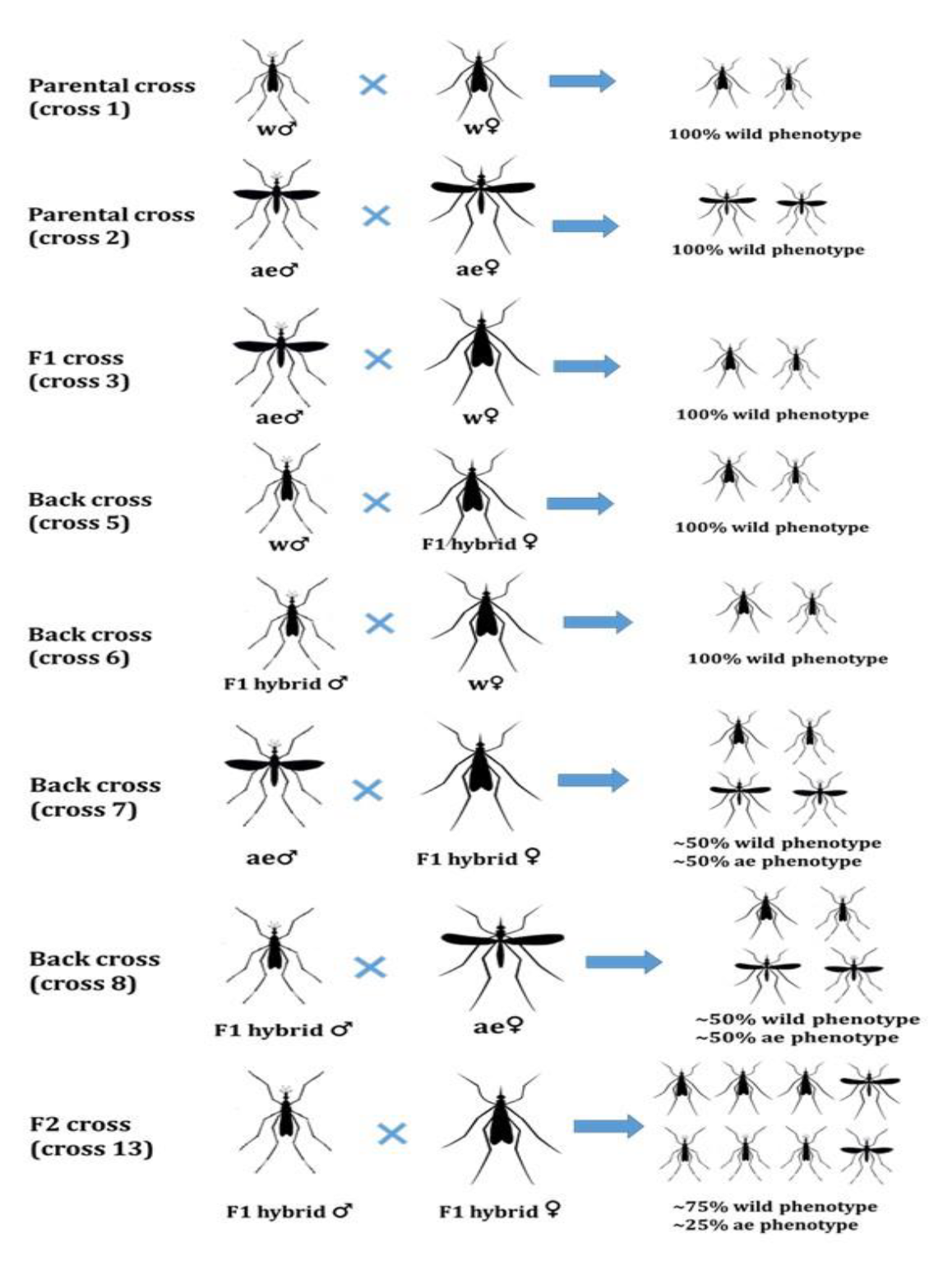
Schematic illustration of different crosses between mutants (*ae*) and wild (w) types.

**Table 1.**
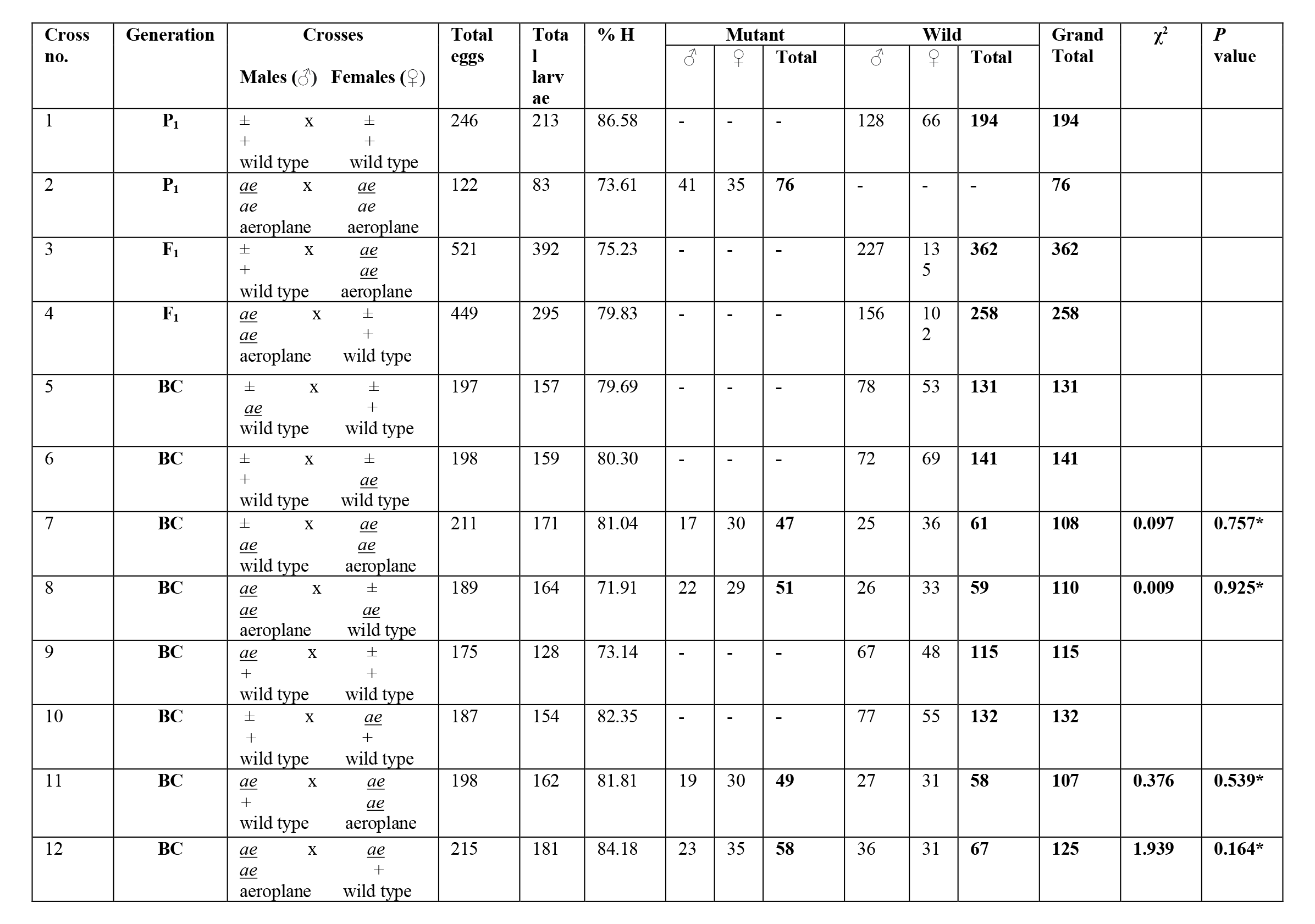
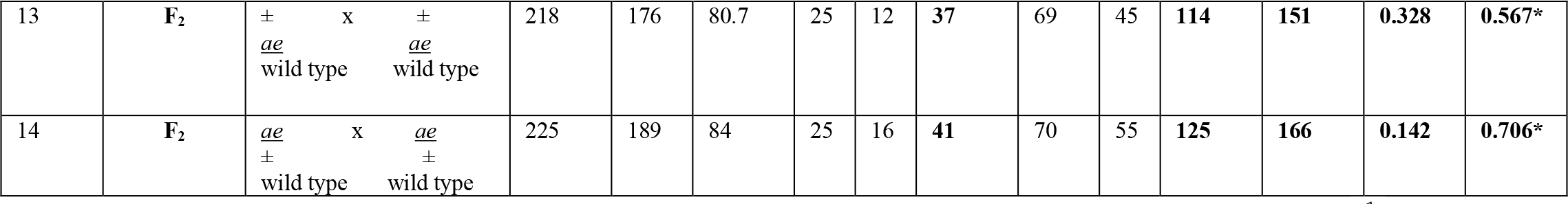
Mode of inheritance of aeroplane mutant (*ae*) in *An. stephensi.* [BC, backcross; **♂**, male; ♀, female. *Statistically non-significant (*P*>0.05)].

Crosses 1 and 2 were performed between males and females of wild-type and *ae* mutants and confirmed the establishment and purity of homozygotes of mutant and wild-type. In the reciprocal crosses (crosses 3 and 4), the F_1_ progeny were wildtype. Therefore, this indicates that *<u>ae</u>* gene is recessive to the wild-type. The absence of the *ae* mutant in the heterogametic males in F_1_ indicates that the gene *<u>ae</u>* is autosomal. Crosses 5, 6, 7, 8 and 13 were derived from the male outcross of the F_1_ progeny (cross 3 of Table 1). Crosses 9, 10, 11, 12 and 14 were derived from the female outcross of the F_1_ progeny (cross 4 of Table 1). When heterozygous F_1_ hybrid progeny were backcrossed to pure-bred wild types (w), no mutant phenotype was observed (crosses 5, 6, 9 and 10 of Table 1); only wild individuals were noticed. However, backcrosses of F_1_ heterozygous progeny with the presumptive parental homozygotes of mutant types resulted both wild and mutant phenotypes (crosses 7, 8, 11 and 12 of Table 1). Results of these backcrosses revealed the approximate 1:1 ratio of wild-type to *ae* mutant. The *χ*^2^ values of these crosses indicate non-significant deviations (for crosses 7 and 8, *χ*^2^ = 0.097, *P* = 0.757 and *χ*^2^ = 0.009, *P* = 0.925; for crosses 11 and 12, *χ*^2^ = 0.376, *P* = 0.539 and *χ*^2^ =1.939, *P* = 0.164, respectively). Some adults of the F_1_ were inbred to yield F_2_ generations (crosses 13 and 14 of Table 1). The mutant and wild type showed 3:1 ratio and no significant *χ*^2^ values were obtained (*χ*^2^ = 0.328, *P* = 0.567 and *χ*^2^ = 0.142, *P* = 0.706). Data from Table 1 clearly demonstrate that *<u>ae</u>* gene is recessive to wild-type, autosomal and its inheritance is monogenic in nature (Table 1).

Results of crosses F_3_ to F_15_ are summarized in Table 2 and data represents the total number of male and female mutants that appeared in each generation. In all filial generations (F_3_–F_15_), the percentage of mutant phenotypes ranged between 15 and 19%. The lowest and highest ratios of mutant and wild mosquitoes were observed in the ratio of 0.16:1 and 0.23:1, respectively. The mean values of fecundity and hatchability per female were ∼60.33 (range 51–73; SEM=6.91±1.78) and ∼57.5 (range 42–71; SEM=9.13±2.36). Data from Table 2 clearly demonstrates that the mutant gene *<u>ae</u>* is transferred over generations with uniform phenotypic expressions.

**Table 2:** The frequency of *ae* mutants over 15 generations after F_2_ cross. (LW, left wing; RW, right wing; DW, double wing; **♂,** male; **♀**, female).

| Gen | Filial | No. Eggs | No. larvae | %H | No. adults | Wild ♂ | Wild ♀ | Total wild (♂+♀) | Mutant ♂ |  |  | Total mutant ♂ | Mutant ♀ |  |  | Total mutant ♀ | Total mutant (♂+♀) | Mutant (%) | Mutant/wild (ratio) | Longevity (day) |
| --- | --- | --- | --- | --- | --- | --- | --- | --- | --- | --- | --- | --- | --- | --- | --- | --- | --- | --- | --- | --- |
|  |  |  |  |  |  |  |  |  | LW | RW | DW |  | LW | RW | DW |  |  |  |  |  |
| G1 | F3 | 131 | 72 | 54.96 | 52 | 26 | 16 | 42 | 2 | 1 | 2 | 5 | 1 | 2 | 2 | 5 | 10 | 19.23 | 0.19:1 | ~30-35 |
| G2 | F4 | 203 | 107 | 57.52 | 78 | 44 | 20 | 64 | 2 | 2 | 1 | 5 | 3 | 4 | 2 | 9 | 14 | 17.94 | 0.21:1 | .. |
| G3 | F5 | 212 | 95 | 44.81 | 80 | 46 | 19 | 65 | 3 | 1 | 2 | 6 | 3 | 4 | 3 | 10 | 15 | 18.75 | 0.23:1 | .. |
| G4 | F6 | 154 | 101 | 65.58 | 76 | 36 | 26 | 62 | 2 | 3 | 2 | 7 | 2 | 2 | 3 | 7 | 14 | 18.42 | 0.22:1 | .. |
| G5 | F7 | 186 | 98 | 52.68 | 68 | 37 | 20 | 57 | 1 | 2 | 1 | 4 | 2 | 3 | 2 | 7 | 11 | 16.17 | 0.19:1 | .. |
| G6 | F8 | 175 | 92 | 52.57 | 79 | 40 | 24 | 64 | 2 | 3 | 2 | 7 | 2 | 4 | 2 | 8 | 15 | 18.98 | 0.23:1 | .. |
| G7 | F9 | 182 | 89 | 48.9 | 77 | 39 | 26 | 65 | 1 | 2 | 2 | 5 | 3 | 3 | 1 | 7 | 12 | 15.58 | 0.18:1 | .. |
| G8 | F10 | 163 | 82 | 50.3 | 71 | 37 | 21 | 58 | 2 | 2 | 1 | 5 | 3 | 2 | 3 | 8 | 13 | 18.30 | 0.22:1 | .. |
| G9 | F11 | 159 | 89 | 55.97 | 73 | 36 | 24 | 60 | 2 | 2 | 2 | 6 | 2 | 3 | 2 | 7 | 13 | 17.80 | 0.21:1 | .. |
| G10 | F12 | 193 | 119 | 61.65 | 79 | 43 | 24 | 67 | 2 | 1 | 2 | 5 | 2 | 3 | 2 | 7 | 12 | 15 | 0.22:1 | .. |
| G11 | F13 | 274 | 166 | 60.58 | 80 | 39 | 26 | 65 | 3 | 3 | 2 | 8 | 2 | 2 | 3 | 7 | 15 | 18.75 | 0.23:1 | .. |
| G12 | F14 | 295 | 172 | 58.3 | 75 | 21 | 23 | 64 | 1 | 2 | 2 | 5 | 1 | 3 | 2 | 6 | 11 | 14.66 | 0.17:1 | .. |
| G13 | F15 | 114 | 81 | 71.05 | 65 | 35 | 20 | 55 | 2 | 1 | 1 | 4 | 3 | 2 | 1 | 6 | 10 | 15.38 | 0.18:1 | .. |
| G14 | F16 | 121 | 84 | 69.42 | 70 | 38 | 22 | 60 | 1 | 2 | 2 | 5 | 1 | 2 | 2 | 5 | 10 | 14.28 | 0.16:1 | .. |
| G15 | F17 | 103 | 73 | 70.87 | 59 | 31 | 19 | 50 | 2 | 2 | 1 | 5 | 2 | 1 | 1 | 4 | 9 | 15.25 | 0.18:1 | .. |

### C. Morphometric analyses of eggs and wings

Morphometric analyses were carried out from eggs and wings of both *ae* mutant and parental T_6_ lines and their comparative significance levels were calculated.

#### a) Egg stage

##### Egg shape

In both the lines, the shape of the eggs was slightly boat shaped in ventral, dorsal and lateral views and black in colour. Anterior and posterior ends were blunt, but sometimes pointed. Ventral surface was concave and dorsal surface was curved (Fig. 3).

**Figure 3:**
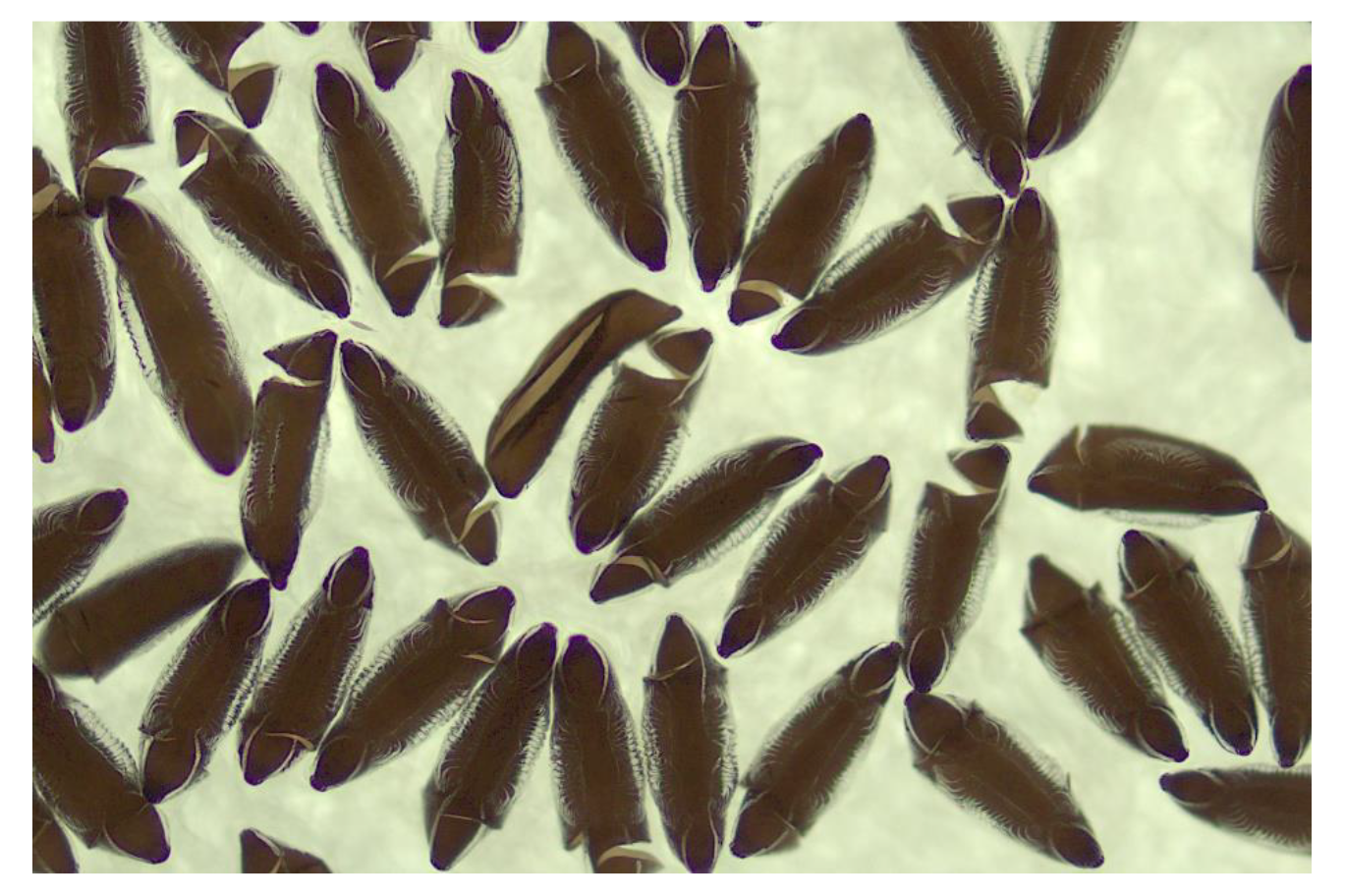
Eggs of *ae* mutant (magnification 10x).

##### Egg size

The maximum egg lengths of DW, RW and LW mutants were 422.5±3.39, 418.5±3.101 and 425±2.564 μm, respectively. The maximum egg widths were 127±1.791, 125±1.539 and 126±1.529 μm for DW, RW and LW mutants, respectively (Table 3). There was no significant difference observed in the egg length and egg width when compared among DW/RW, DW/LW and RW/LW mutants (*P*>0.05) (Table 4).

**Table 3:** Egg measurement of mutant strains. ER (egg-float ridge), EL (length of egg), EW (width of egg), EFL (length of egg-float), EFW (width of egg-float) among *ae* mutant eggs.

| Parameter | DW |  | RW |  | LW |  |
| --- | --- | --- | --- | --- | --- | --- |
|  | M±SEM | SD | M±SEM | SD | M±SEM | SD |
| ER | 15.52±0.159 | 0.696 | 15.31±0.109 | 0.477 | 15.42±0.139 | 0.607 |
| EL | 422.5±3.39 | 15.174 | 418.5±3.101 | 13.869 | 425±2.564 | 11.47 |
| EW | 127±1.791 | 8.013 | 125±1.539 | 6.882 | 126±1.529 | 6.805 |
| EFL | 208±1.716 | 7.677 | 207±1.791 | 8.013 | 206.5±1.817 | 8.127 |
| EFW | 69.5±1.697 | 7.591 | 67.5±1.758 | 7.864 | 70±1.622 | 7.254 |

**Table 4:** Comparative significance values of length and width among DW, RW and LW mutant eggs.

|  |  | DW/RW | DW/LW | RW/LW |
| --- | --- | --- | --- | --- |
| Length (µm) | Mean | 420.5 | 423.75 | 421.75 |
|  | F value | 1.2 | 1.75 | 1.46 |
|  | P value | 0.347 | 0.1158 | 0.2085 |
| Width (µm) | Mean | 125.5 | 126.5 | 124.75 |
|  | F value | 1.86 | 1.39 | 1.34 |
|  | P value | 0.0926 | 0.2398 | 0.2648 |

##### Egg float

Floats were present on both sides of the egg surface, which is filled with air that helps in floating in water. The maximum egg-float lengths of DW, RW and LW mutants were 208±1.716, 207±1.791 and 126±1.529 μm, respectively. The maximum-egg float widths were 69.5±1.697, 67.5±1.758 and 70±1.622 μm of DW, RW and LW mutants, respectively (Table 3).

##### Egg-float ridge

The ridges on the float present on both lateral surface on eggs serves as an important identifying character for different biological forms in *Anopheles* species. They were short and closer to ventral than dorsal surface. For DW, RW and LW mutants the number of float ridges was observed 15.52±0.159 (range 15–17), 15.31±0.109 (range 15–16) and 15.42±0.139 (range 15–17), respectively (Table 3). No significant difference was observed in different parameters of egg measurement of DW when compared with its parental T_6_ isofemale lines (*P*>0.05) (Table 5).

**Table 5:** Comparative significance values between DW mutant and parental (T_6_) eggs.

|  | Egg length | Egg width | Egg-float length | Egg-float width |
| --- | --- | --- | --- | --- |
| Mean | 420 | 129.333 | 213 | 69.333 |
| F value | 1.51 | 1.03 | 18.95 | 1 |
| P value | 0.2252 | 0.47834 | <0.0001 | 0.5 |

### b) Wing measurements

In DW mutant male, the range of wing length is 2.635–3.081 mm and width 0.471–0.591 mm; whereas in female, the range of wing length is 2.5–3.656 and width 0.603–0.852 mm (Table 6). In RW male, the range of wing length and width are 2.539–2.862 and 0.537–0.581 mm; in female wings, length and width are 2.893–3.467 and 0.727–0.846 mm, respectively. In LW male, the range of wing length and width are 2.546–3.055 and 0.445–0.505 mm. In LW female, wing length and width are 3.164–3.539 and 0.744–0.852 mm, respectively (Fig. 4a and b). In parental T_6_ males, the range of wing length and width are 3.1–3.2 and 0.55–0.69 mm; while in parental T_6_ females, the range of wing length and width are 3.1–3.4 and 0.726–0.841 mm, respectively (Table 7; Fig. 4c and d). No significant difference was observed in wing measurement when compared between the mutant and parental T_6_ isofemale lines (*P*>0.05) (Tables 8 and 9).

**Figure 4:**
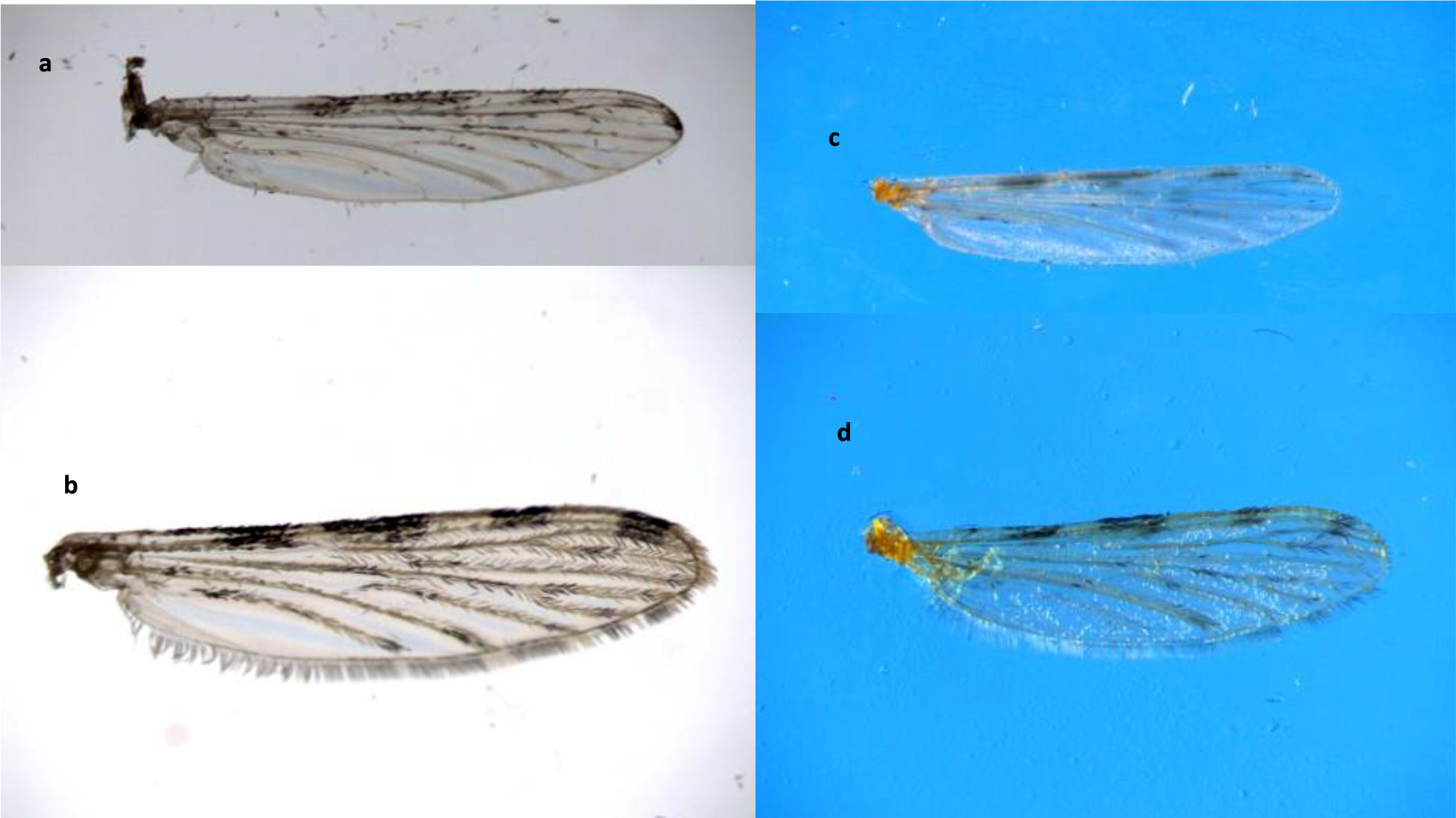
Male (a) and female (b) wings of T_6_ Isofemale parent and male (c), female (d) wings of *ae* mutant lines.

**Table 6:** Wing measurements of *ae* mutant.

| Mutant | Male |  |  | Female |  |
| --- | --- | --- | --- | --- | --- |
|  | Length (mm) | Width (mm) |  | Length (mm) | Width (mm) |
| **LW** |  |  |  |  |  |
| Range | 2.546-3.055 | 0.445-0.505 |  | 3.164-3.539 | 0.744-0.852 |
| M±SEM | 2.71±0.089 | 0.518±0.023 |  | 3.43±0.068 | 0.792±0.0194 |
| SD | 0.2007 | 0.053 |  | 0.152 | 0.0434 |
| **RW** |  |  |  |  |  |
| Range | 2.539-2.862 | 0.537-0.581 |  | 2.893-3.467 | 0.727-0.846 |
| M±SEM | 2.726±0.057 | 0.563±0.008 |  | 3.163±0.121 | 0.786±0.022 |
| SD | 0.129 | 0.019 |  | 0.270 | 0.049 |
| **DW** |  |  |  |  |  |
| Range | 2.635-3.081 | 0.471-0.591 |  | 2.5-3.656 | 0.603-0.852 |
| M±SEM | 2.816±0.068 | 0.527±0.018 |  | 2.991±0.144 | 0.737±0.029 |
| SD | 0.167 | 0.044 |  | 0.408 | 0.083 |

**Table 7:** Wing measurements of T_6_ paternal isofemale lines.

| **T<sub>6</sub> Isofemale** |  |  |  |  |
| --- | --- | --- | --- | --- |
|  | **Male** |  | **Female** |  |
| **Wing measurement** | Length (mm) | Width (mm) | Length (mm) | Width (mm) |
| **Range** | 3.1-3.2 | 0.55-0.69 | 3.1-3.4 | 0.726-0.841 |
| **M±SEM** | 3.150±0.016 | 0.631±0.023 | 3.359±0.03 | 0.783±0.015 |
| **SD** | 0.041 | 0.057 | 0.093 | 0.038 |

**Table 8:** Comparative statistical analyses between T_6_ parental isofemale and *ae* mutant wings.

| **Mosquito strains** | **Male** |  |  |  | **Female** |  |  |  |
| --- | --- | --- | --- | --- | --- | --- | --- | --- |
|  | **Length** |  | **Width** |  | **Length** |  | **Width** |  |
|  | **M±SEM** | **SD** | **M±SEM** | **SD** | **M±SEM** | **SD** | **M±SEM** | **SD** |
| **T<sub>6</sub> Isofemale** | 3.150±0.017 | 0.0407 | 0.631±0.024 | 0.0576 | 3.359±0.038 | 0.093 | 0.783±0.015 | 0.039 |
| ***Ae* mutant** |  |  |  |  |  |  |  |  |
| **DW** | 2.816±0.068 | 0.167 | 0.528±0.018 | 0.044 | 2.991±0.144 | 0.408 | 0.737±0.029 | 0.084 |
| **RW** | 2.726±0.057 | 0.129 | 0.563±0.008 | 0.019 | 3.164±0.121 | 0.270 | 0.786±0.022 | 0.049 |
| **LW** | 2.71±0.089 | 0.201 | 0.518±0.023 | 0.053 | 3.43±0.068 | 0.152 | 0.792±0.019 | 0.043 |
| Non-significant | *P*>0.05 |  | *P*>0.05 |  | *P*>0.05 |  | *P*>0.05 |  |

**Table 9:** Comparative analysis of *P* and F values of length and width of males and females between T_6_ and *ae* mutants [non-significant, NS (*P*>0.05)].

| **Mosquito strains** | **Male** |  |  |  | **Female** |  |  |  |
| --- | --- | --- | --- | --- | --- | --- | --- | --- |
|  | **Length** |  | **Width** |  | **Length** |  | **Width** |  |
|  | **F value** | ***P* value** | **F value** | ***P* value** | **F value** | ***P* value** | **F value** | ***P* value** |
| ***Ae* mutant wing** |  |  |  |  |  |  |  |  |
| **DW/T<sub>6</sub>** | 14.8 | 0.003 | 1.85 | 0.221 | 20.19 | 0.002 | 4.9 | 0.049 |
| **RW/T<sub>6</sub>** | 9.74 | 0.014 | 9.57 | 0.024 | 8.08 | 0.021 | 1.55 | 0.317 |
| **LW/T<sub>6</sub>** | 23.39 | 0.001 | 1.22 | 0.436 | 2.57 | 0.164 | 1.2 | 0.413 |

### D. Cytogenetic studies of ae mutant and T_6_ isofemale parental line – 3Li inversion

In the present study, heterozygous paracentric inversions, *i*/+ were observed on 3L arm of *ae* mutants. The percent of the inversion recorded is 19.64% (Table 10). The tentative breakpoints of the inversions on 3L arm involved 42A–44C (Fig. 5a and b). A similar 3L*i* inversion was also observed at a higher percentage (47.45%) in its T_6_ parental isofemale line.

**Table 10:** Percent chromosomal inversions in *ae* mutant and parental T_6_ isofemale strains of *An. stephensi*.

| Strain | Types of chromosome inversions | Chromosome arm involved | Tentative break points | % inversion | No. of positive slides/total slides prepared |
| --- | --- | --- | --- | --- | --- |
| *ae* mutant | Heterozygous | 3L (*i/+*) | 42A–44C | 19.64% | 11/56 |
| Parental Iso T<sub>6</sub> | Heterozygous | 3L (*i/+*) | 42A– 44C | 47.45% | 28/59 |

**Figure 5:**
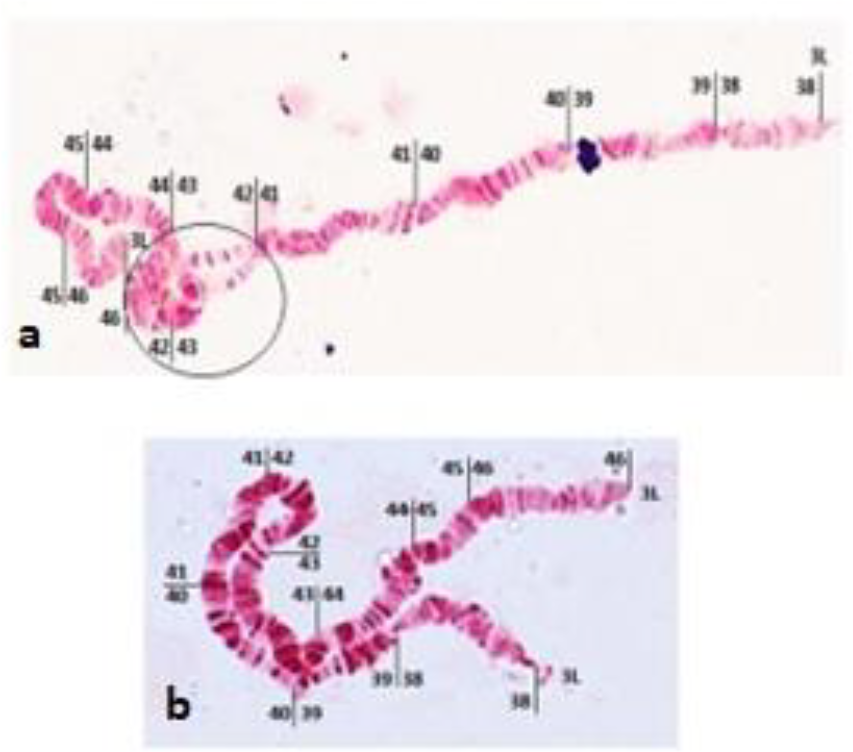
Photomap of polytene chromosome from ovarian nurse cells: (a) *ae* mutant showing an inversion, and (b) standard 3L arm without inversion from T_6_ Iso lines of *An. stephensi*.

## Discussion

Establishment and characterization of phenotypic markers have great applications in expanding the genetic knowledge of non-model insect pests and vectors. These markers can be used in devising specific crosses, constructing genetic maps, and identifying loci of quantitative and/or qualitative traits. The phenotypic marker genes can be used as guides for insect transformation studies and to bridge classical and applied research. The study isolates and characterizes a novel phenotypic open winged mutant in *An. stephensi*. Genetic and cytogenetic studies of *An. stephensi* continue to be an important area of research for understanding the biology, behaviour and ecology to develop better, more specific, and eco-friendly means of controlling mosquito vectors.

The mutant mosquito is termed as *aeroplane* based on its abnormal wing posture. It has straightened out wings on either one or both sides with a bent abdomen, which looks exactly like an aeroplane. The bent abdomen is observed in both the sexes of males and females, especially in DW mutants. The positions of the wing in these three types of mutants (DW, LW and RW) are unique. In the typical form of DW, both wings are widely extended forming 90° angle with the longitudinal axis of the body. In LW and RW, the wings are extended towards the left and right sides of the body, respectively, while the other wings lie on the body same like wild type. The wings are of the same shape and texture as those of wild type. The wings of mutants are flat, and the venation is similar like that of wild type. There are no marked differences observed in wing shape, structure and the halteres are also similar in shape to those of the wild type. All the mutants are strong, active, fertile and comparable to their wild counterpart. They are seen flying like normal mosquitoes inside the cages and remain alive for 30–40 days. They are competent to take blood meal both from artificial blood feeding systems (Haemotek ®) and a mouse host as well. Over generations, good fecundity and hatchability were observed, 75–90 and 80%, respectively (Table 2). The open wing character is prominently visible soon after eclosion. The mutant strains are currently being maintained in TIGS insectary for more than 25 generations.

Genetic analysis demonstrated that the *<u>ae</u>* gene is autosomal and recessive and monogenic in nature. Crossing of F_3_ to F_15_ progeny confirms the stability of the mutant gene in the mutant population and the wing character is unique. Precise morphometric analysis of mutant eggs shows the egg-float ridge numbers ranged between 15 and 16, which is helpful in its classification under intermediate variants. The 3L*i* inversion in *ae* mutant persists as in its parental line (T_6_), however with a reduced frequency (19.64% in *ae* mutants and 47.45%, in Iso T_6_ lines) [25]. Notably, inversion is the most prominent adaptation mechanism for new biotic and abiotic environment. Often this inversion could be developed as a specific fixed marker for that species [26]. Inversions are physical rearrangements of a chromosome segment, which limit the exchange of variation between two alternate pairs of alleles. These diverged adaptive gene complexes function as supergenes in many organisms. Chromosomal inversions are widespread and have been extensively associated with latitudinal and longitudinal patterns among members of the *Anopheles* species [24]. Sixteen paracentric inversions in the chromosomes of *An. stephensi* have been reported by visualizing loop formation in chromosomes from large homozygous/heterozygous inversions [27, 28].

Recent advances in gene editing CRISPR/Cas9 to develop mutant mosquitoes heavily rely upon expression of phenotypic markers for confirming the transformation events. There have been reports of morphologically distinct mutants developed through radiation, where the phenotypes can be easily visualized in larval and adult stages. The larval colour mutants in *An. stephensi* and their genetic basis of inheritance like stripe, controlled by a codominant gene [9]; grey, an autosomal recessive gene [16]; greenish brown, an autosomal recessive gene; red eye, a recessive sex-linked gene [29]; diamond palpus, an autosomal recessive gene [30]; golden-yellow, an autosomal recessive gene [31]; white eye, a recessive sex-linked gene; yellow body larvae, an autosomal recessive gene [15]; greyish black, an autosomal recessive gene [32]; ruby-eye <u>(*ru*),</u> an autosomal recessive gene [10]; and dark mutant <u>(*da*),</u> an autosomal recessive gene [33] have been reported by several pioneer workers [34]. The tests for allelism among certain larval colour mutants have also been reported earlier in *An. stephensi*. The mutant larvae brown (*<u>br</u>*) is allelic to green (*<u>gr</u>*) [35], and grey (*<u>gy</u>*) is allelic to greenish black (*<u>gbl</u>*) [16]. The linkage relationship between the hairless antenna (*<u>hla</u>*) and ruby eye (*<u>ru</u>*) mutant revealed the suppression of *<u>hla</u>* by the *<u>ru</u>* gene in heterozygous condition [10]. The green thorax (*<u>gt</u>*) allele is recessive and autosomal to wild-type. The linkage studies showed no linkage between *<u>ru</u>* and *<u>gt</u>* [19]. Among the adults, there are easily visualized mutant phenotypes in the wing venation, eye, and antenna in *Culex quinquefasciatus* and *Culex fatigans*, respectively [36, 37]. Similarly, gene knockout studies targeting the flight muscle gene have been conducted in *Ae. aegypti* making the transformed flightless mosquitoes, phenotypically distinct from the other non-transformed mosquitoes [38]. This flightless gene was used as a phenotypic marker in the development of CRISPR-based precision-guided sterile insect technique (pgSIT) in *Ae. aegypti*. We propose that the *ae* mutant described in this study could be an excellent marker of *An. stephensi* as it expresses with complete penetrance and high viability in its adult stages. The mutant could also be used extensively for conducting basic genetic experiments in *An. stephensi*. We hypothesize that the novel *ae* mutant phenotype in *An. stephensi* is like the aeroplane wing mutant previously reported in *Drosophila melanogaster* [39]. In *Drosophila*, when mutants were crossed with wild types, F_1_ progeny gave wild-type offspring only, but in F_2_ progeny the wing character reappeared in both sexes [40]. Similar to our findings based on the experimental crosses, the gene of wing mutant in *Drosophila* was also recessive, autosomal and present on the 2nd chromosome. It would be interesting to do molecular characterization of the *<u>ae</u>* mutant gene and know the possible reasons of the open winged phenotype in *An. stephensi*.

## Conclusion

Aeroplane, a new wing mutant in *An. stephensi* is reported for the first time. The genetic analyses of the above mutant demonstrate that the mutant gene, *<u>ae</u>* is autosomal, recessive and non-sex-linked, and might be controlled by a single gene. Three types of wings orientations are observed in *ae* mutants, i.e., DW, LW and RW and no significant differences were observed in the egg and wing morphometrics when compared with the parental line. Cytogenetic study confirmed the persistence of the 3L*i* inversion in *ae* mutants like their parental lines. Further molecular genetics study is necessary, which would enlighten several novel avenues to identify target gene/s for causing mutations. It would also advance our knowledge of the evolution of mutation in vectors which might consider the effective ways to manage vector control.

## Ethics statement

Biosafety approval for mosquito maintenance facility (approval Ref. No. TIGS 3rd IBSC March 2018) and Institutional ethical approval for use of human blood for mosquito feeding were obtained [approval Ref. No. inStem/IEC-12/002, Bangalore from Institute for Stem Cell Science and Regenerative Medicine (inStem)]. The blood used during feeding mosquitoes for colony maintenance was collected from a licensed human blood bank (Lion’s Blood Bank, Bangalore, India).

## Abbreviations

ae: aeroplane
T_2_: TIGS-2
T_6_: TIGS-6
DW: double wing
LW: left wing
RW: right wing

## Acknowledgements

The authors thank Dr. Rakesh Mishra, Director, Tata Institute for Genetics and Society (TIGS), Bangalore, India for his support and guidance. We are thankful to Dr. Susanta Kumar Ghosh, Former Scientist G and Head, ICMR-National Institute of Malaria Research, Bangalore, India for guiding the establishment of TIGS insectary. We are grateful to Mr. Joydeep Roy for his technical support. Animal work in the NCBS/inStem Animal Care and Resource Center was partially supported by the National Mouse Research Resource (NaMoR) grant #BT/PR5981/MED/31/181/2012;2013-2016 & 102/IFD/SAN/5003/2017-2018 from the Department of Biotechnology, GoI. The authors thank Tata Institute for Genetics and Society (TIGS) India for funding researchers involved in this work.

## Author contributions

CG: Wrote the manuscript; isolated, established and maintenance of aeroplane (*ae*) mutant line; designed and performed crossbreeding experiments; morphometric analyses of eggs; data recording and statistical analyses; and inversions in polytene chromosomes.

SM: Insectary-related work, isofemale line, mutant line maintenance and wing dissections.

NK: Insectary-related work and imaging of the mutants.

CKR: Insectary-related work, wing measurements and photograph.

SGJ: Insectary-related work and wing measurements.

SK: Schematic diagram, imaging of mutant and normal mosquitoes and helping in reviewing the manuscript draft.

SS: Reviewing the manuscript and intellectual inputs.

SunS: Coordinating and overseeing the entire insect work and helping in reviewing the manuscript draft.

All authors read and approved the final manuscript.

